# A meta-analysis of bulk RNA-seq datasets identifies potential biomarkers and repurposable therapeutics against Alzheimer’s disease

**DOI:** 10.1101/2023.09.17.558173

**Authors:** Anika Bushra Lamisa, Ishtiaque Ahammad, Arittra Bhattacharjee, Mohammad Uzzal Hossain, Ahmed Ishtiaque, Zeshan Mahmud Chowdhury, Keshob Chandra Das, Md Salimullah, Chaman Ara Keya

**Author notes:** These authors have contributed equally (Anika Bushra Lamisa, Ishtiaque Ahammad, Arittra Bhattacharjee).

## Abstract

Alzheimer’s disease (AD) poses a major challenge due to its impact on the elderly population and the lack of effective early diagnosis and treatment options. In an effort to address this issue, a study focused on identifying potential biomarkers and therapeutic agents for AD was carried out. Using RNA-Seq data from AD patients and healthy individuals, 12 differentially expressed genes (DEGs) were identified, with 9 expressing upregulation (*ISG15, HRNR, MTATP8P1, MTCO3P12, DTHD1, DCX, ST8SIA2, NNAT,* and *PCDH11Y*) and 3 expressing downregulation (*LTF, XIST,* and *TTR*). Among them, *TTR* exhibited the lowest gene expression profile. Interestingly, functional analysis tied *TTR* to amyloid fiber formation and neutrophil degranulation through enrichment analysis. These findings suggested the potential of *TTR* as a diagnostic biomarker for AD. Additionally, druggability analysis revealed that the FDA-approved drug Levothyroxine might be effective against the Transthyretin protein encoded by the *TTR* gene. Molecular docking and dynamics simulation studies of Levothyroxine and Transthyretin suggested that this drug could be repurposed to treat AD. However, additional studies using in vitro and in vivo models are necessary before these findings can be applied in clinical applications.

## Introduction

Alzheimer’s disease (AD) is a degenerative neurological condition characterized by a progressive and irreversible decline in cognitive and functional abilities, resulting in significant impairment in routine tasks and social interactions [1]. Various environmental and genetic risk factors have been linked with its onset [2], [3]. The pathological mechanisms of the disease are mostly distinguished by the formation of amyloid plaques (Aβ plaques), neurofibrillary tangles, and neuronal degeneration in the brain [4]. Furthermore, Alzheimer’s disease (AD) is a matter of significant public health concern on a global scale, both in the United States and in several other countries around the world [5], [6]. Furthermore, it is noteworthy that this condition ranks as the fifth most prevalent cause of death in the elderly population of the United States. Additionally, it is estimated that approximately 35 million individuals globally are impacted by this disease, with estimations indicating that this figure will increase to 65 million by the year 2030 [7]. Additionally, the number of cases of Alzheimer’s disease in Asia, in developed as well as developing countries, is affected by age, gender, and cultural factors [8], [9], [10]. In Bangladesh, there is a lack of precise epidemiological data on the number of AD patients, and awareness of the disease is still in the stages of development. The lack of awareness has resulted in ongoing difficulties for patients and their families. Moreover, limited funding for AD research makes it difficult for a lower-middle-income country like Bangladesh to effectively manage the disease. Therefore, it is time to consider the disease and its management proactively and take the necessary measures [11].

Several studies have been conducted in recent times regarding the etiology of AD, with the majority of the research indicating that the cause of AD was genetic in nature [12]. Despite the increasing number of genes that have been suggested to impact vulnerability to Alzheimer’s disease (AD), the comprehension of their mechanisms and the enhancement of disease management are still restricted by challenges in understanding the functional implications of genetic associations [13], [14]. Recent research has demonstrated that various mechanisms of gene expression regulation, including interactions between mRNA and transcription factors, non-coding RNAs, alternative splicing, and variants, may have an impact on the process of neurodegeneration and an increasing number of developments are being observed in the concurrent analysis of transcriptome data, to investigate the consequences of recently identified genetic risk factors on the transcriptome level [15].

The area of molecular biology and genomics has grown tremendously in the previous decade, with the most recent development content including “omics” technologies such as genomics, proteomics, transcriptomics, and metabolomics [16]. The transcriptomic concept refers to the complete set of transcripts found in various cell types, tissues, or organs. This encompasses both coding and noncoding RNA molecules that are responsible for encoding proteins [17]. Several studies have utilized the transcriptomics approach to identify biomarkers that differentiate individuals with Alzheimer’s disease from those without Alzheimer’s. The identification of differentially expressed genes (DEGs) through RNA-Seq analysis is an essential part of the study of biological pathways implicated in various neurological disorders. The purpose of conducting Differential Expression Gene (DEG) analysis is to identify genes that exhibit potential overexpression or underexpression in the context of a disease state, relative to a control group that remains unaffected [18]. Dysregulation of gene expression, whether it be overexpression or underexpression, can lead to disruptions in various biological pathways such as metabolic and immune pathways, which eventually result in the development of diseases [19]. Differentially expressed genes (DEGs) may exert an impact on the initiation of neurodegenerative disorders like Alzheimer’s disease. Furthermore, it is plausible that there may exist divergences in the gene expression profiles across distinct regions of the brain [20]. Determining whether the transcriptional alterations may yield cumulative impacts on established disease susceptibility factors and disease-associated pathways is of paramount significance [21]. The incidence of Alzheimer’s disease (AD) may exhibit an upward trend with advancing age, and anomalous transcriptional alterations give rise to pathogenic mechanisms associated with the disease [22]. The application of differentially expressed genes (DEGs) in systemic biology studies can identify significant functional components and central hub genes that are linked to the development of various diseases. This approach may involve the use of various tools such as Gene Ontology (GO) [23] and Kyoto Encyclopedia of Genes and Genomes (KEGG) [24] biological pathways.

The goal of the study was to identify differentially expressed genes from bulk RNA-Seq data and suggest potential biomarkers as well as find any repurposable drug candidates, if available. For this purpose, a number of analyses such as differential gene expression, druggability, molecular docking and dynamics simulation were to be carried out.

## Methods

### Retrieval of NGS data

The RNA-Seq datasets of 211 patients with Alzheimer’s (AD=133) and non-Alzheimer’s (control=78), were obtained in FASTQ format from the public database Gene Expression Omnibus (GEO) (https://www.ncbi.nlm.nih.gov/geo/) (Project ID: PRJNA675355, PRJNA767074, PRJNA232669, PRJNA796229, PRJNA516886, PRJNA279526, PRJNA413568, PRJNA675355) in the National Center of Biotechnology Information (NCBI) website. These samples were subsequently processed using various computational tools. These steps have been represented through a flowchart (**Figure 1**).

**Figure 1:**
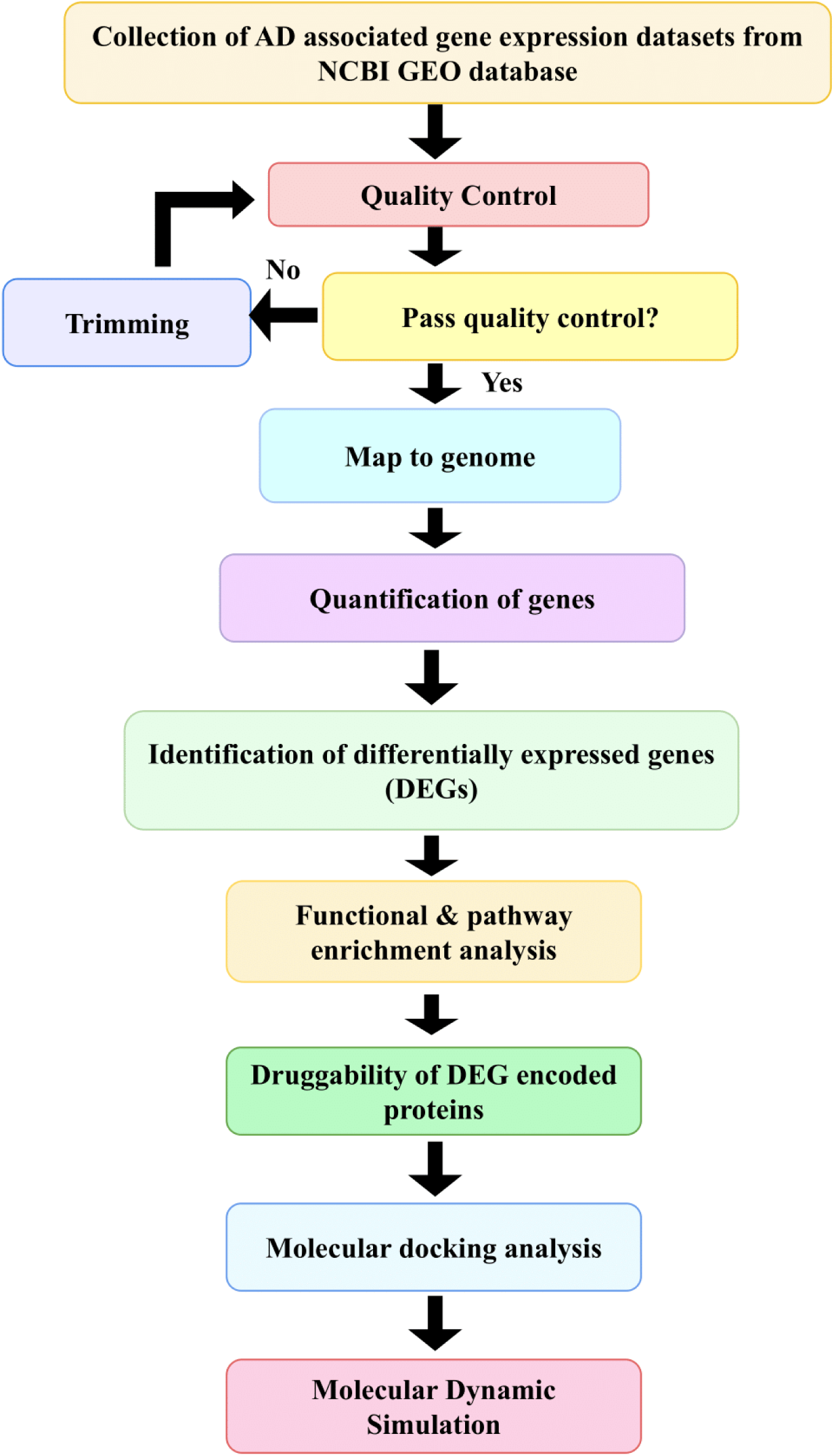
Overall workflow of the study.

### Processing and alignment of the reads

FastQC, an open-source software, was employed to evaluate the quality of the raw reads (https://www.bioinformatics.babraham.ac.uk/projects/fastqc/). HISAT2 was used to align all the raw reads to the reference genome of *Homo sapiens* GRCh38.p13 (GCA_000001405.28) from Ensembl [25]. Afterwards, the mapped reads were then assigned to Ensembl genomic features defined in GRCm38.69 (http://primerseq.sourceforge.net/gtf.html). The number of reads per gene were quantified with the use of FeatureCounts (http://subread.sourceforge.net/).

### Differential gene expression analysis

The quantification of RNA-Seq relies on read counts that are assigned to genes in a probabilistic manner. For the minimization of batch effect, ComBat-seq function of sva package was used for each project [26]. In order to compute differential expression, the statistical approach DESeq2 was used to predict DEGs, and false discovery rate (FDR) was used to correct p-values and identify true DEGs. The fold change (FC) of each gene was calculated between control and AD groups, and genes with p-adjusted value < 0.05 and |Log2FC| > 1.45 were considered significant DEGs [27].

### Functional and pathway enrichment analysis of DEGs

To understand the biological functions and pathways associated with the differentially expressed genes (DEGs), enrichment analyses of Gene Ontology (GO) and KEGG pathways were performed. Enrichment analysis was carried out using Enrichr (https://maayanlab.cloud/Enrichr/), which is a web-based tool for functional enrichment analysis. GO terms with a corrected p-value ≤0.05 were considered significantly enriched. For pathway enrichment analysis, the KEGG database was used in the Enrichr web server and pathways with a p-value ≤0.05 were considered significantly enriched.

### Druggability of DEG encoded proteins

The protein sequences of DEGs were retrieved from UniProtKB (https://www.uniprot.org/help/uniprotkb) and queried in the DrugBank database (https://go.drugbank.com/) as drug targets. Proteins were considered potential druggable targets if their sequences had a high degree of similarity (E value-<10^-100^, bit score>100) to those in the DrugBank database. Those that had no hits were regarded as novel targets [28].

### Molecular docking

Chemical structure of Levothyroxine was obtained from DrugBank. It was docked against the target downregulated transthyretin protein encoded by *TTR* gene using Webina 1.0.3 web server (https://durrantlab.pitt.edu/webina/). Following successful molecular docking, the protein-ligand interactions were visualized using Pymol (https://pymol.org/2/) and Poseview(https://proteins.plus/).

### Molecular dynamic simulation

100 ns Molecular dynamics (MD) simulation was carried out using the GROningen MAchine for Chemical Simulations aka GROMACS (version 2020.6) for the apo *TTR* and Levothyroxine-*TTR* complex [29]. The proteins were embedded in the TIP3 water model [30], [31]. The whole system was energetically minimized using CHARMM36m force-field [32]. The systems were neutralized using K^+^ and Cl^-^ ions. Following energy minimization, isothermal-isochoric (NVT) equilibration, and Isobaric (NPT) equilibration of the system were executed. Afterward, 100 ns production MD simulation was run. Using MD simulation data, root mean square deviation (RMSD), root mean square fluctuation (RMSF), radius of gyration (Rg), and solvent accessible surface area (SASA), and Hydrogen bond analysis were conducted. The ggplot2 package (https://ggplot2.tidyverse.org/) in RStudio was utilized for generating the graphs for each of these analyses. All MD simulations were performed in the high-performance simulation stations running on Ubuntu 20.04.4 LTS operating system located at the Bioinformatics Division, National Institute of Biotechnology, Bangladesh.

## Results

### Quantification of the high quality raw reads

FastQC v0.11.5 was utilized to check the quality of the raw sequencing data obtained from NCBI. A total of 211 raw sequences were evaluated and all of them were found to be of high quality. Following the alignment of the reads to the human reference genome, a total of 62,702 genes were identified which were forwarded to DEG analysis in the quantification step.

### Differential gene expression analysis revealed significantly upregulated and downregurated genes

A total of 10730 differentially expressed genes (DEGs) were identified in AD patient samples, with 7814 genes being upregulated and 2916 genes being downregulated. These DEGs were visualized using Rstudio, and various plots such as MA plots and volcano plots were constructed. In Figure 2, the MA plot illustrates the DEGs, with the upregulated DEGs shown on the top and the downregulated DEGs on the bottom. The significant DEGs were marked by blue dots. In Figure 3, the volcano plot displayed the DEGs, with the upregulated DEGs located on the right and the downregulated DEGs on the left. The significant DEGs (p value < 0.05) are indicated by the yellow color while the rest was shown in red.

**Figure 2:**
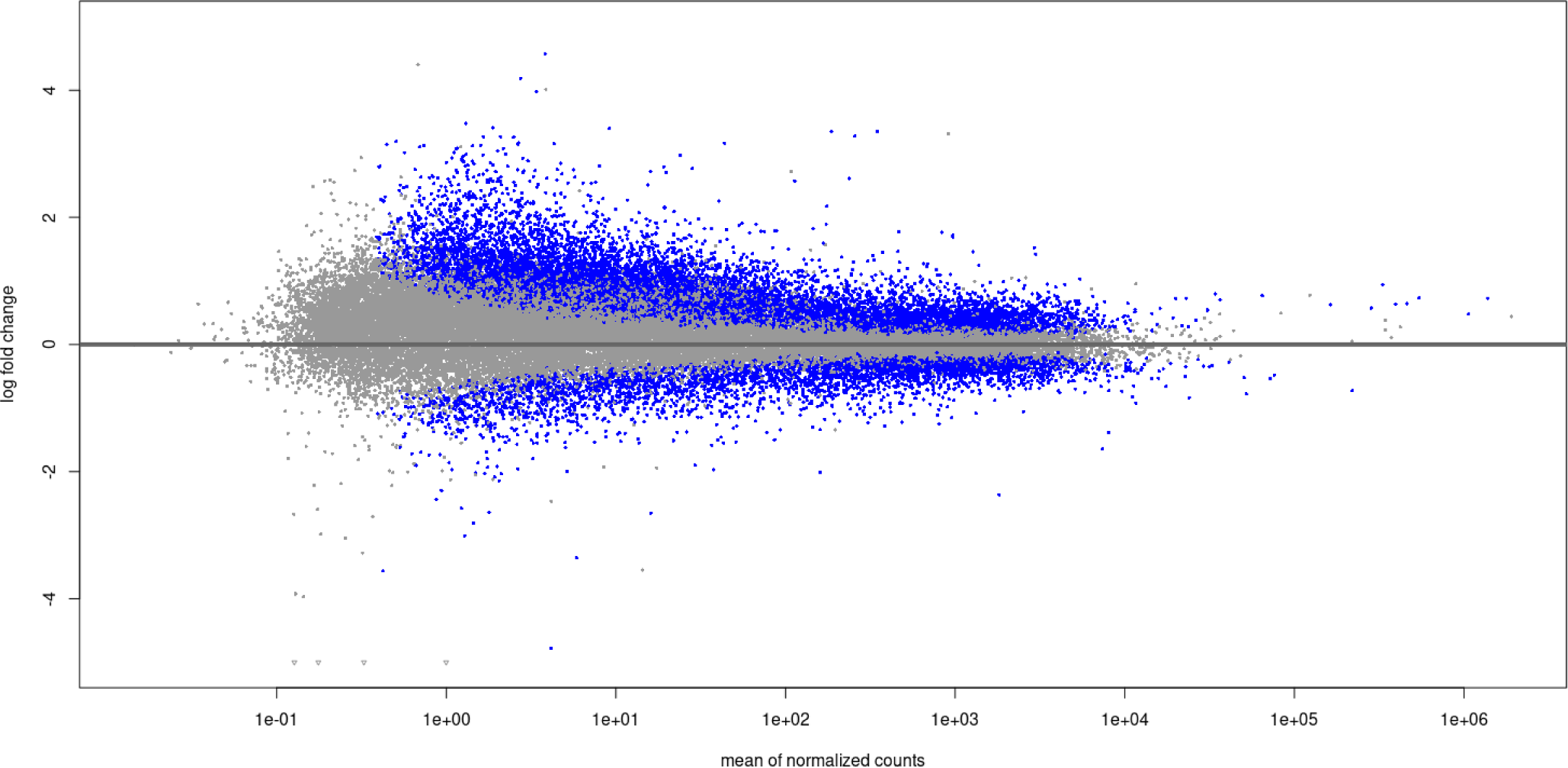
MA plot of differentially expressed genes identified in AD patient samples. The upregulated and downregulated genes were displayed at the top and bottom of the plot respectively. The significant DEGs were indicated by the blue dots.

**Figure 3:**
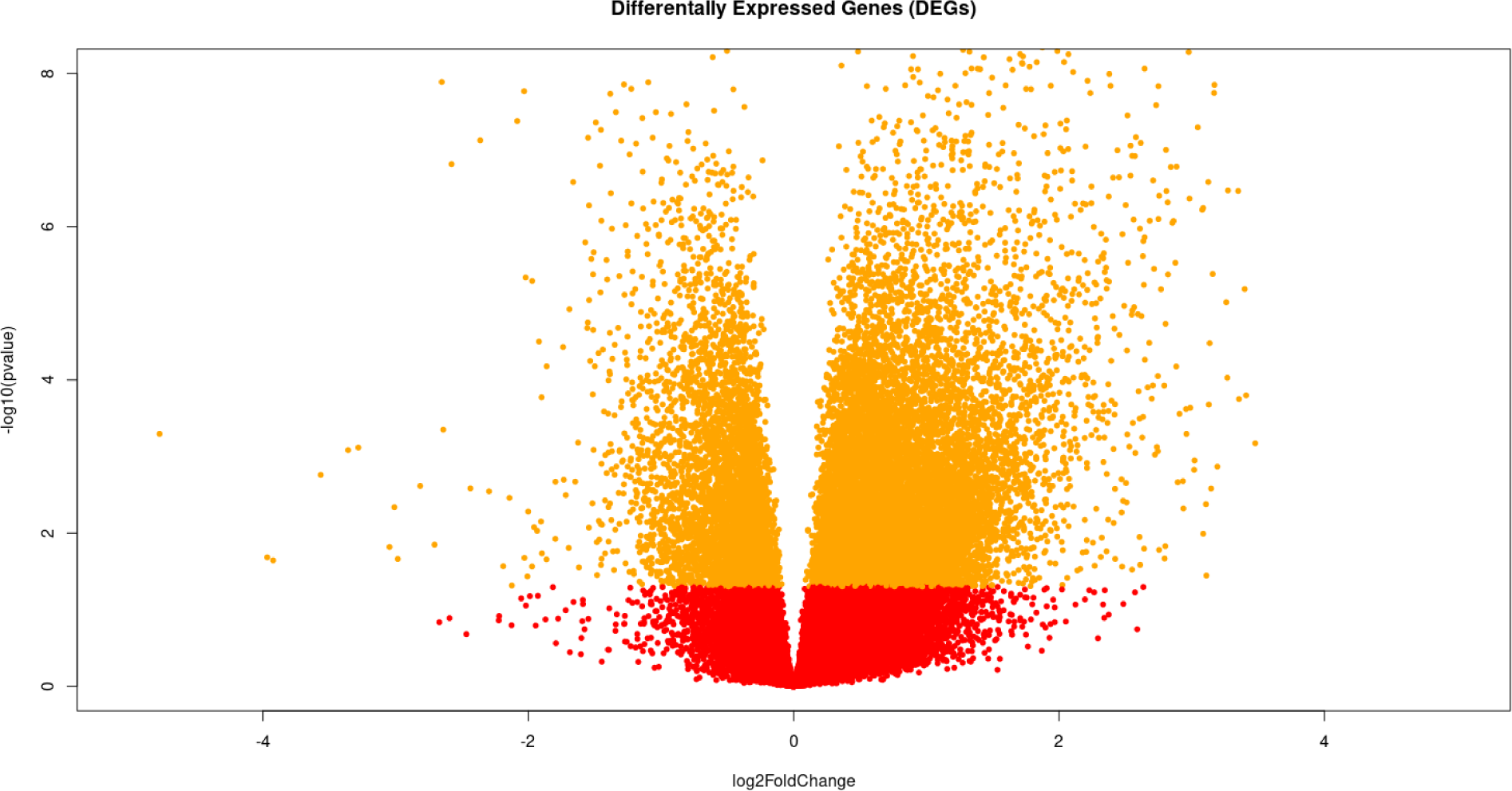
Volcano plot of differentially expressed genes. The upregulated and downregulated genes were depicted on the right and left respectively. The genes were indicated using yellow dots.

### Significant functional and pathway enrichment analysis of DEGs

**Table 1** demonstrated that out of 10730 DEGs, a total 12 genes passed the cutoff value (|Log2FC| > 1.45) for significance. Among these 12 genes, 9 DEGs were upregulated and 3 DEGs were downregulated. This figure also depicted that in Alzheimer’s disease, *PCDH11Y* was the most upregulated (log2foldchange value = 1.889662998) and *TTR* was the most downregulated (log2foldchange value = -2.361971992) DEGs. The GO functional and KEGG pathway enrichment analysis of these 12 DEGs showed that 11 GO-Biological Process (GO-BP) terms, 9 GO-Cellular Component (GO-CC) terms, 6 GO-Molecular Function (GO-MF) terms, 1 KEGG pathway and 5 Reactome pathways were associated with the 2 downregulated genes, respectively (**Supplementary Table 1**). The downregulated gene-associated KEGG pathway was thyroid hormone synthesis and Reactome pathways were Amyloid fiber formation, metal sequestration by antimicrobial proteins, neutrophil degranulation and innate immune systems (Figure 4a). Moreover, 4 GO-Biological Process (GO-BP) terms, 2 GO-Cellular Component (GO-CC) terms, 2 GO-Molecular Function (GO-MF) terms, 1 KEGG pathway were associated with the 9 upregulated genes. Among them, one upregulated gene *ISG15* was found to be involved in RIG-I-like receptor signaling pathway (Figure 4b).

**Table 1:**
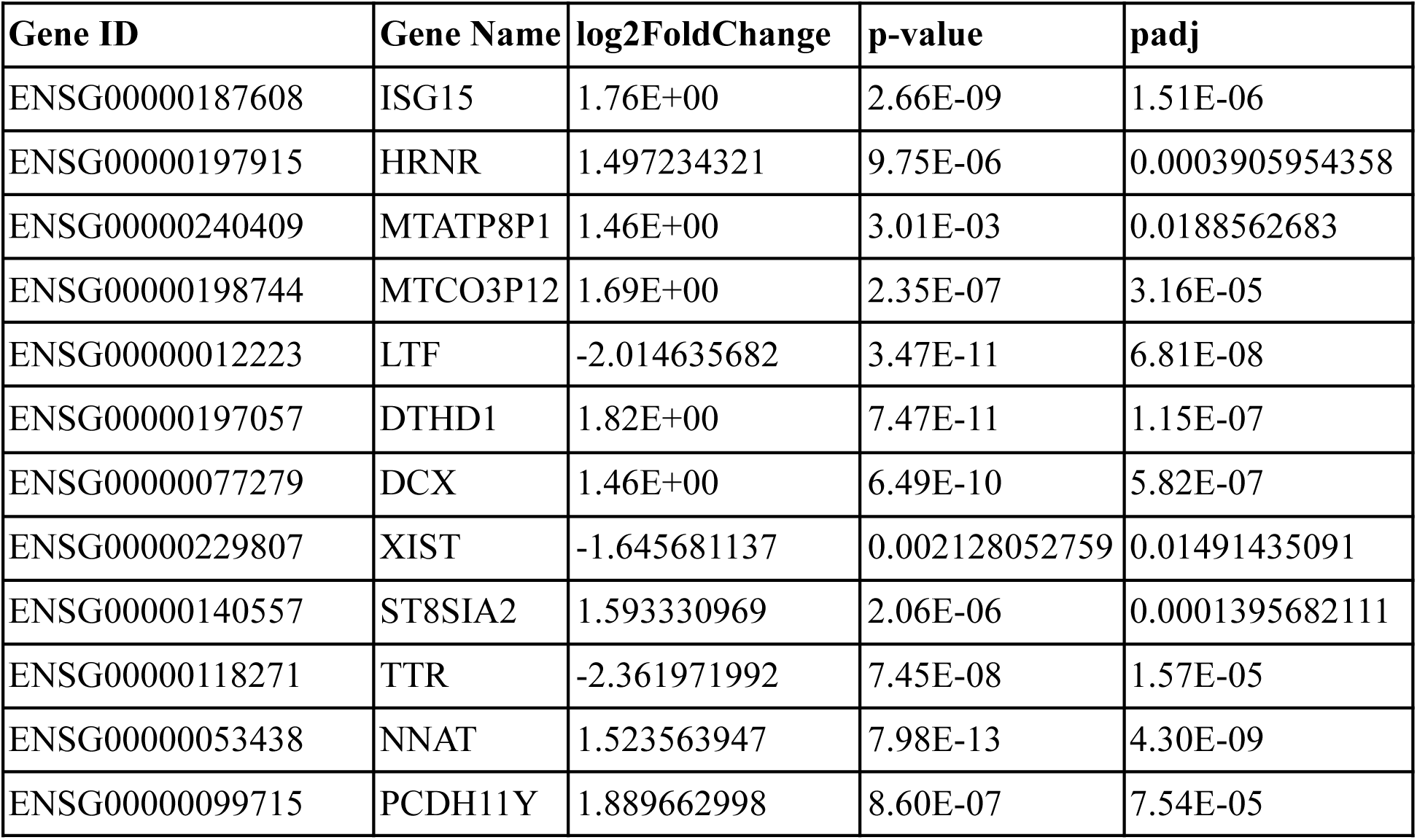
Significant upregulated and downregulated genes.

**Figure 4.**
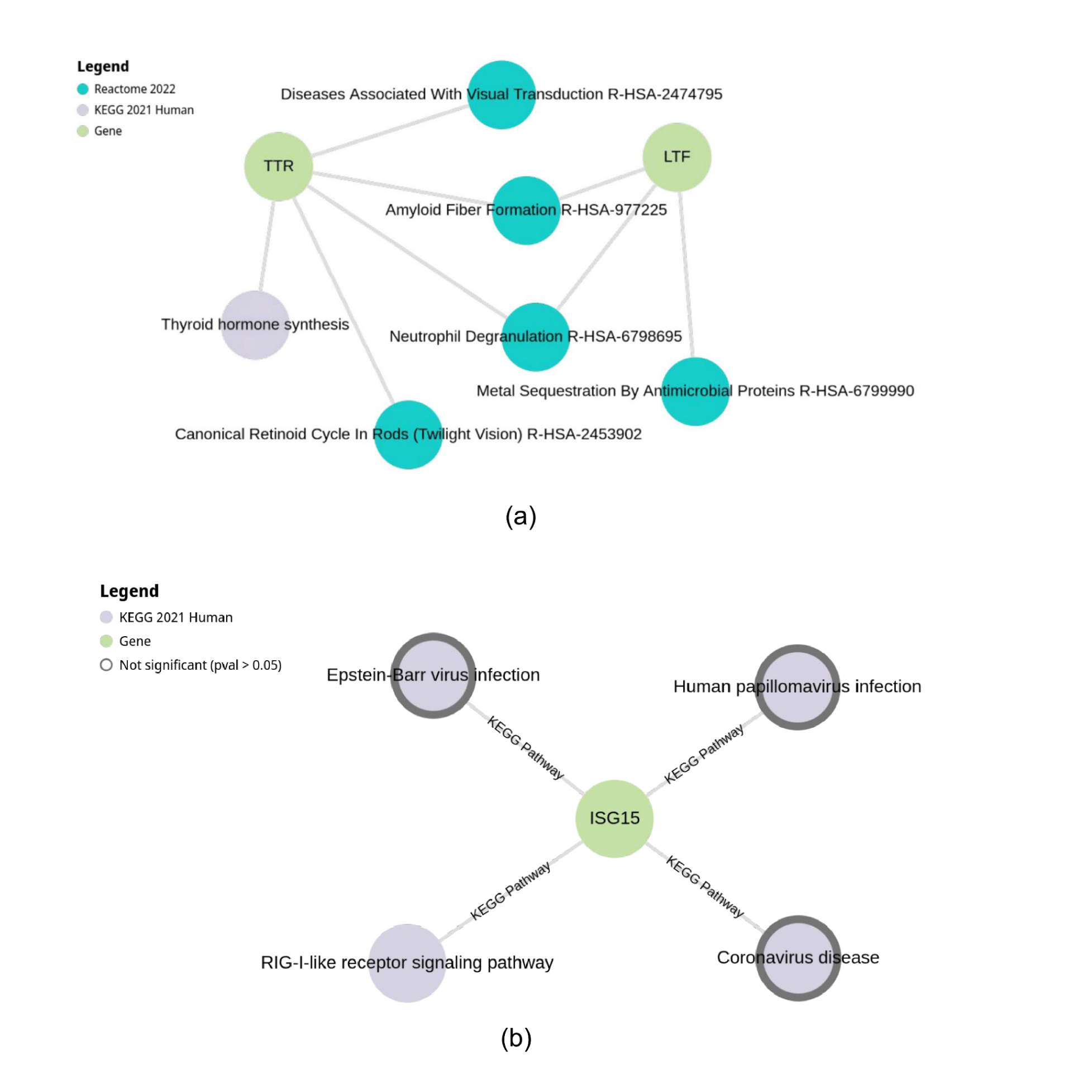
**(a):** Significantly enriched pathway of the downregulated genes by using KEGG and Reactome database; **(b)** Significantly enriched pathway of the upregulated genes by using KEGG database.

### Levothyroxine was identified as a potential drug against TTR gene product Transthyretin

The DrugBank webserver was used to find potential drugs that might target the 4 downregulated genes. It revealed that only one gene (*TTR*) had a corresponding FDA-approved drug called Levothyroxine. The remaining genes did not match to any drugs in the DrugBank database, suggesting that they can be considered as potential novel therapeutic targets.

### Molecular docking analysis revealed molecular interactions between the drug and target protein

Utilizing Webina 1.0.3 web server, a molecular docking analysis had been carried out between the FDA-approved drug Levothyroxine and the predicted structures of the *TTR* encoded protein called Transthyretin. The molecular interactions between the ligand Levothyroxine and Transthyretin indicated a significant binding energy value of -5.1 kcal/mol. According to docking analysis, *TTR* gene interacted with Levothyroxine through Arg103A, Asp99A, Thr119A, Ala120A, Ser100A. (Figure 5)

**Figure 5:**
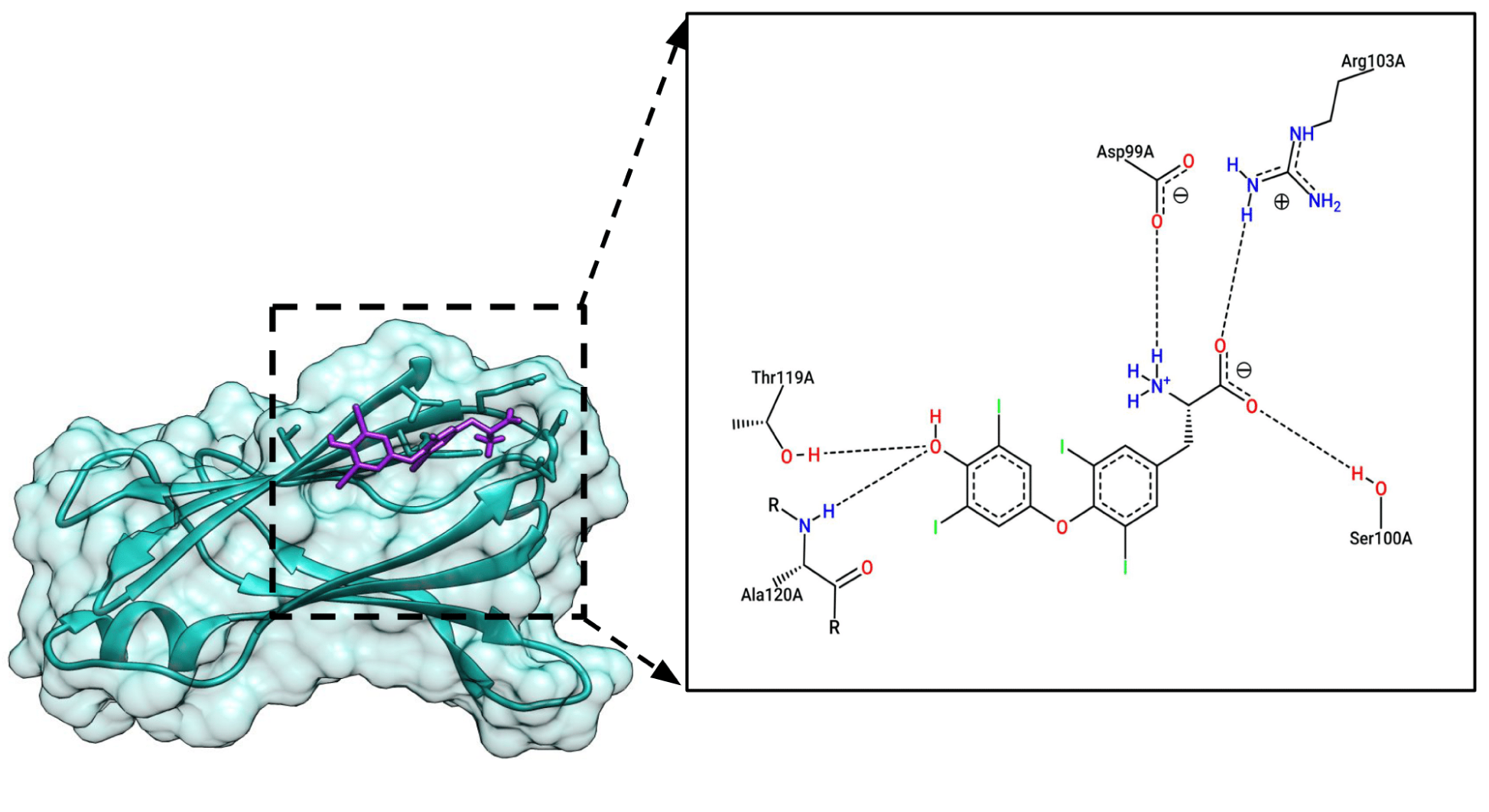
Molecular Docking between the *TTR* and Levothyroxine. *TTR* gene interacts with Levothyroxine through Arg103A, Asp99A, Thr119A, Ala120A, Ser100A.

### MD simulation analysis confirmed the stability of drug-receptor complex

Root Mean Square Deviation (RMSD) calculation was carried out in order to assess stability of the systems. Change in RMSD value corresponds to conformational changes of the protein as a result of ligand binding. The purple line represents the RMSD profile of apo receptor whereas the green line depicts drug-receptor complex (Figure 6). After 50ns, the RMSD value of the drug-receptor complex did not increase beyond ∼2.5 nm whereas the apo receptor RMSD value gradually increased up to ∼4.0 nm. Root Mean Square Fluctuation (RMSF) analysis was utilized to determine the regional flexibility of the protein. The higher the RMSF, the higher is the flexibility of a given amino acid position. Figure 7 demonstrates the RMSF profile of apo receptor and drug-receptor complex. The major RMSF peaks were observed at around the 20^th^ residue and the 85^th^ residue. In the peak near the 85^th^ residue, the apo receptor showed higher mobility. The radius of gyration is a measure to determine its degree of compactness. A relatively steady value of radius of gyration means stable folding of a protein. Fluctuation of radius of gyration implies the unfolding of the protein. According to Figure 8, the Levothyroxine-receptor complex went under less folding according to the Rg (nm) values. Solvent Accessible Surface Area (SASA) is used in MD simulations to predict the hydrophobic core stability of proteins. The higher the SASA value, the higher the chance of destabilization of the protein due to solvent accessibility. The SASA value of apo receptor and drug-receptor was ∼65 nm^2^ initially. However, after 90 ns, the values of both proteins overlapped (Figure 9).

**Figure 6:**
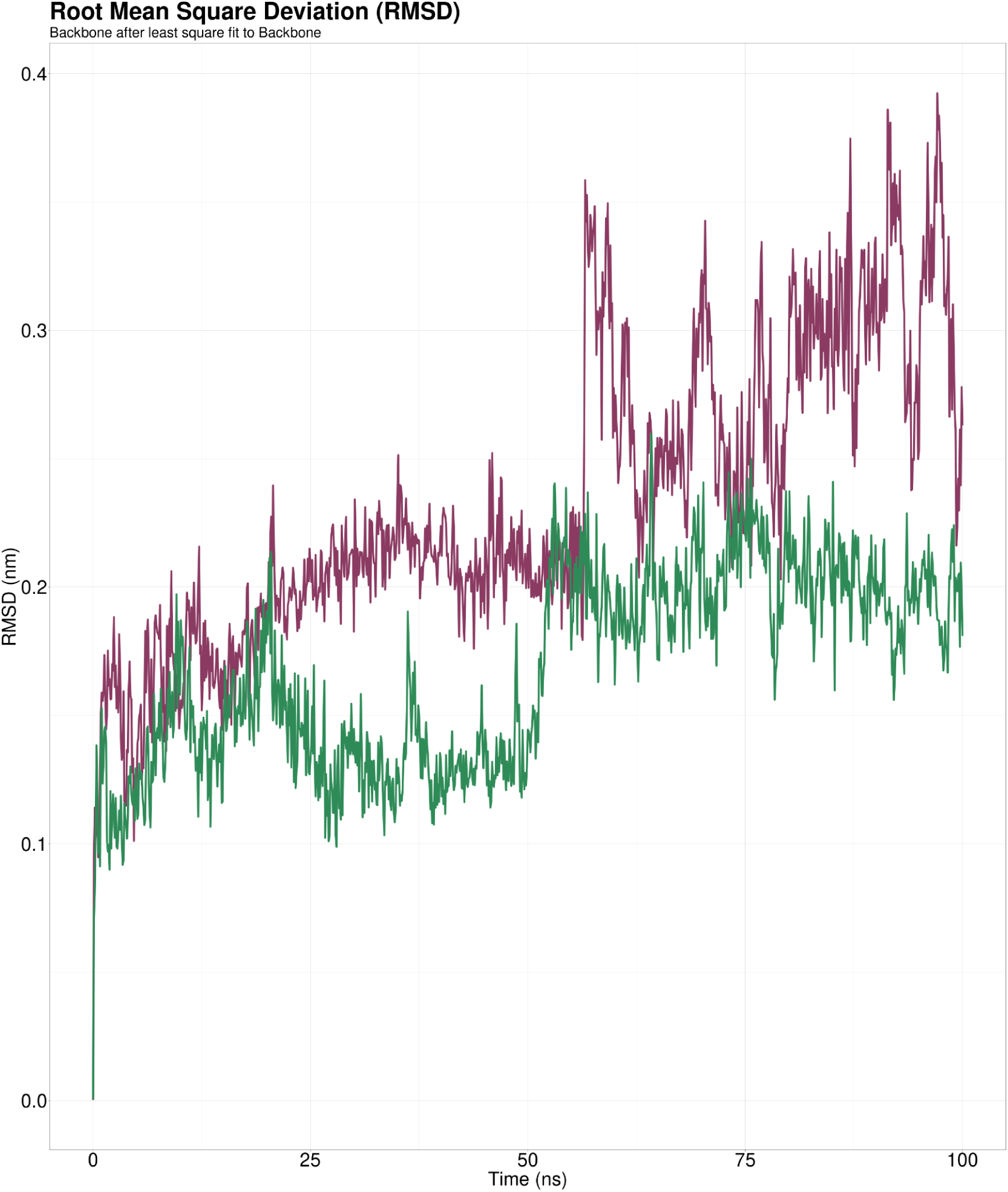
RMSD profile of apo receptor (Purple) and Levothyroxine-receptor complex (Green).

**Figure 7:**
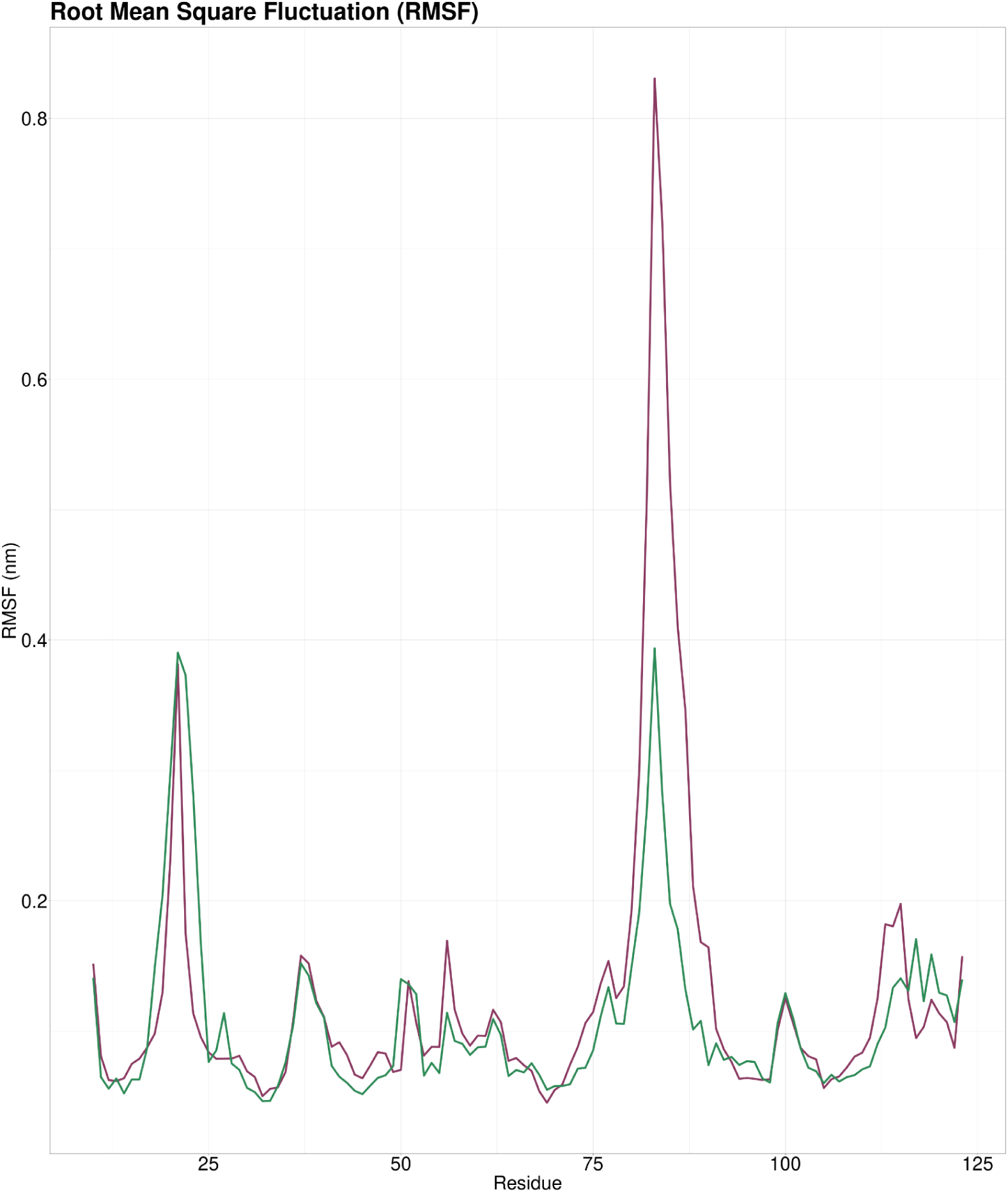
RMSF profile of apo receptor (Purple) and Levothyroxine-receptor complex (Green).

**Figure 8:**
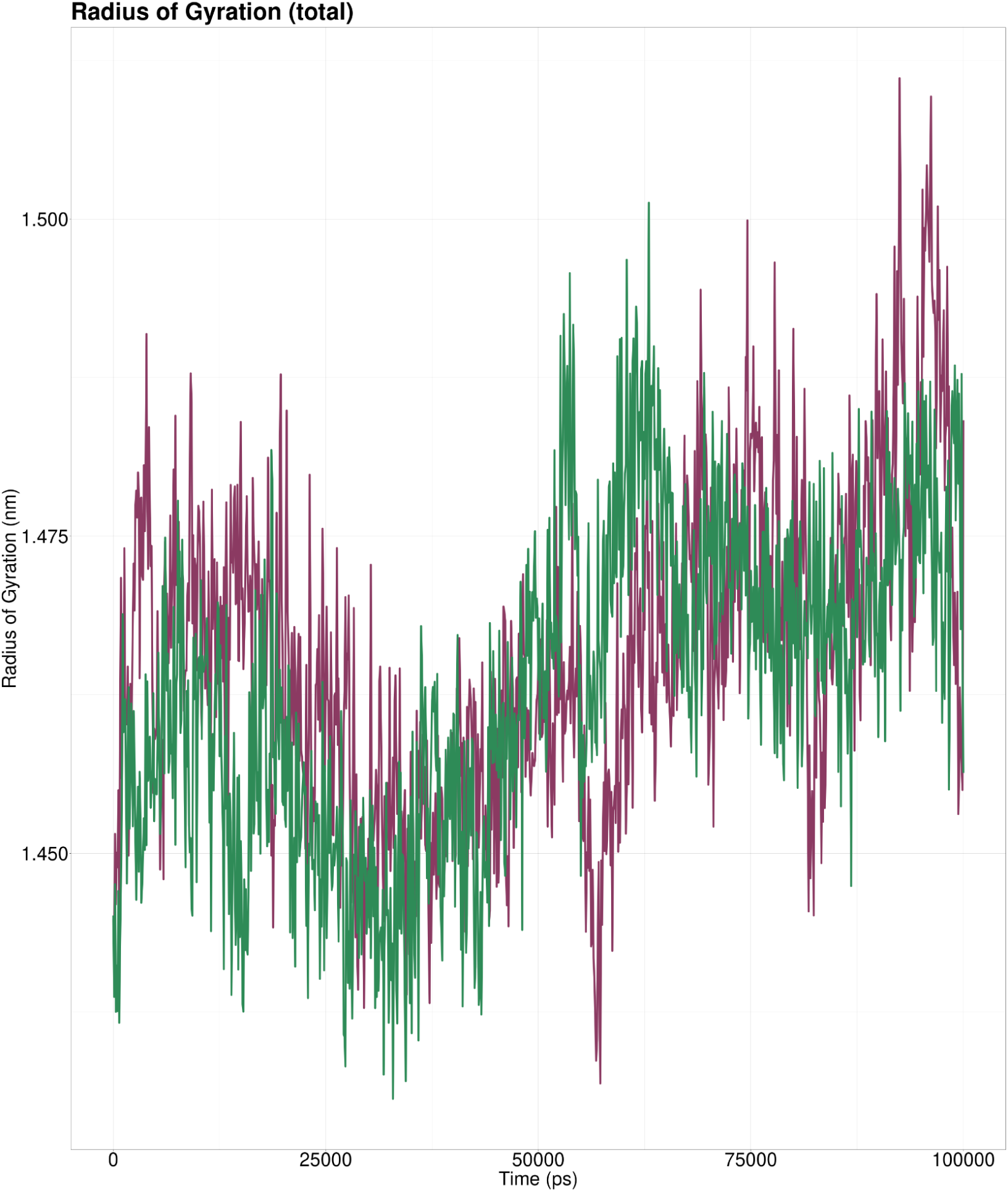
Rg profile of apo receptor (Purple) and Levothyroxine-receptor complex (Green).

**Figure 9:**
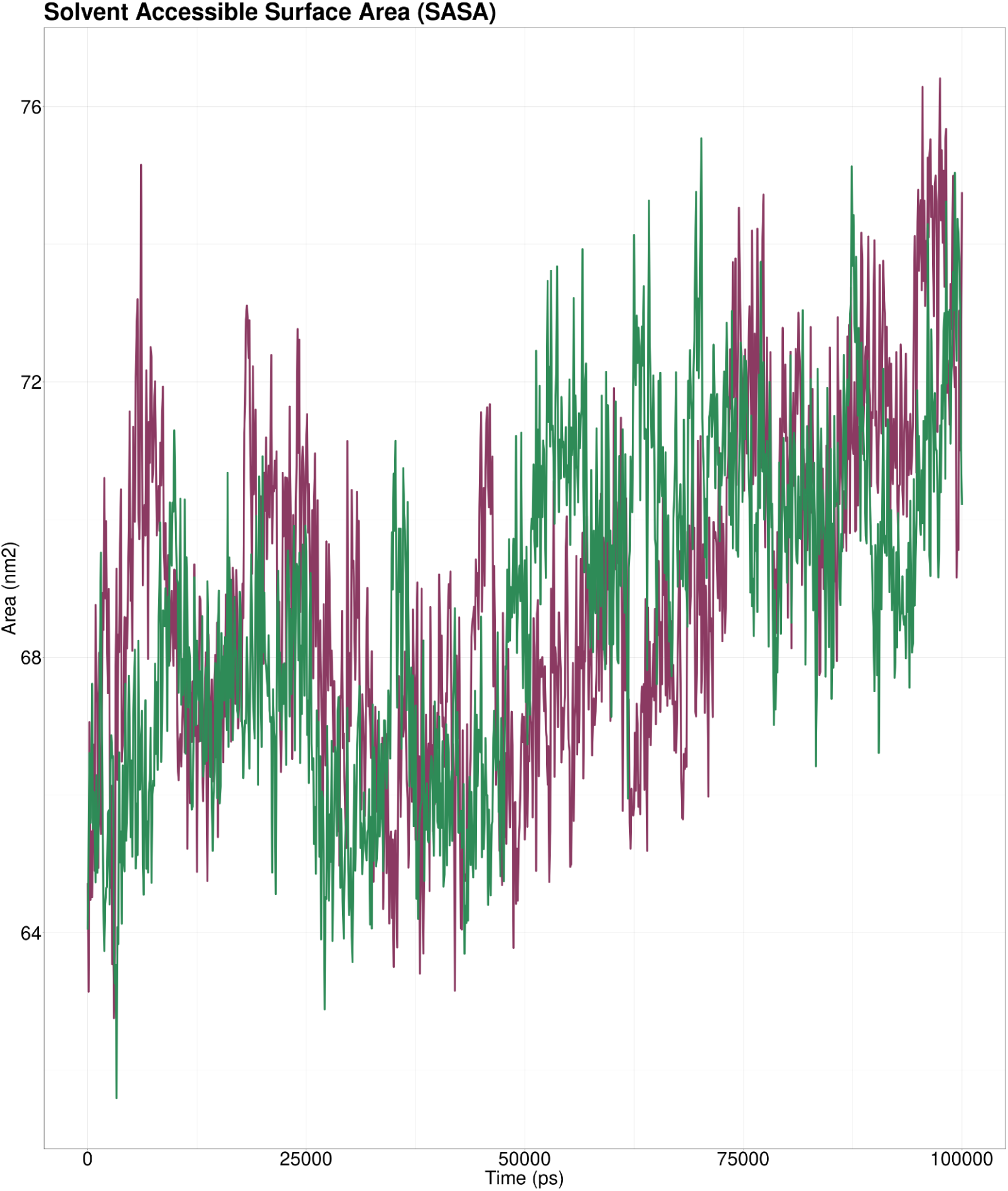
SASA profile of apo receptor (Purple) and Levothyroxine-receptor complex (Green).

## Discussion

AD is a progressive and intricate condition that affects multiple brain functions. Besides the causative genes, various risk factors can also contribute to the progression of the disease. Therefore, using transcriptomics analysis is essential in comprehending the underlying mechanisms of the disease and identifying potential targets for treatment.

In this study, 211 RNA-Seq samples were collected from the GEO database and these samples were found to have high-quality scores. This score is an indication of the accuracy of the base call, and a higher score is generally preferable. However, it is common for the quality of reads to decrease towards the 3’ end, and if the quality of certain bases falls below a certain threshold, they need to be removed. For this reason, the trimming process is necessary to eliminate poor-quality bases, trim adaptor sequences, and ensure high-quality reads. Here, the trimming step was not required since the sample reads were already of high quality. Differentially expressed genes (DEGs) play a crucial role in identifying the biological pathways involved in various diseases including NDs. The main objective of DEG analysis is to identify genes that are either upregulated or downregulated in a disease state compared to healthy controls. Because the upregulation or downregulation of specific genes can cause disruptions in metabolic, immune, and other pathways, potentially leading to disease development [33]. Therefore, identifying DEGs can help us understand the mechanisms underlying disease and develop targeted therapies to treat them. Here, *ISG15, HRNR, MTATP8P1, MTCO3P12, DTHD1, DCX, ST8SIA2, NNAT, PCDH11Y* were found to be most significantly upregulated while the most significantly downregulated genes included *LTF, XIST,* and *TTR*. GO functional analysis revealed that the Alzheimer’s-causing genes were significantly enriched with positive regulation of osteoblast proliferation and development, bone morphogenesis, purine-containing compound metabolic process, regulation of Tumor Necrosis Factor superfamily cytokine production, positive regulation of Toll-Like Receptor 4 signaling pathway, negative regulation of ATP-dependent activity which causes bone loss, inflammation, damage to nerve cells and surrounding brain tissue which were previously reported by other studies as well [34], [35], [36], [37], [38], [39]. One CC term phagocytic vesicle was significantly associated with AD. This was observed to hinder phagocytic process and can cause reduced clearance of accumulated dystrophic neurites [40]. In accordance with other studies, we have also found MF terms such as Cysteine-Type endopeptidase inhibitor activity, Iron ion binding, protein Serine/Threonine kinase activator activity, protein kinase activator activity to be enriched in downregulated genes [41], [42], [43]. Finally, one upregulated gene named *ISG15* was predicted to be involved in RIG-I-like receptor signaling pathway which may lead to the inflammatory response through the activation of type I IFN production in AD [44]. Additionally, two downregulated genes such as *TTR* and *LTF* in AD were predicted to be involved in thyroid hormone synthesis, amyloid fiber formation, diseases associated with visual transduction, neutrophil degranulation, and canonical retinoid cycle in rods (Twilight Vision) pathways. The downregulation of these two genes may lead to the cognitive impairment, amyloid-beta accumulation and retinal dysfunction and visual impairments due to the disruption in thyroid hormone signaling [45], [46], [47].

*TTR* gene product Transthyretin has been reported to inhibit Amyloid-β accumulation and thus prevent the spread of AD [48], [49]. Hyperactivation of neutrophils is a common feature associated with progression of AD. Our analysis uncovered a relationship between neutrophil degranulation and the *TTR* gene as well [50], [51]. Since we have also identified *TTR* as the most downregulated gene in AD patients, it can be predicted as a potential biomarker for AD. Previous studies have also suggested that this gene can be targeted for treating AD [52]. However, the novelty of our study lies in the fact that following whole transcriptome analysis, we looked for the druggability of the targets and discovered that the FDA-approved drug, Levothyroxine, might be an effective repurposable drug against the Transthyretin protein encoded by the *TTR* gene. In this study, Transthyretin protein and Levothyroxine drug showed effective interaction with a high binding energy (−5.1 kcal/mol) which indicated that Levothyroxine can potentially activate the proteins activity. The stability of a Levothyroxine-receptor complex was investigated in this study using molecular dynamics simulations. In order to assess the conformational changes in the protein structure upon drug binding, Root Mean Square Deviation (RMSD) analysis was performed. The drug’s steady binding to the receptor was shown by the gradual increase in the apo receptor’s RMSD profile around 4.0 nm and the drug-receptor complex’s RMSD value being stable at 2.5 nm after 50 ns. The regional flexibility of the protein was ascertained using the Root Mean Square Fluctuation (RMSF) technique. The ∼20^th^ and ∼85^th^ residues were where the RMSF peaks were found. When compared to the drug-receptor complex, the apo receptor had more mobility at the peak near the 85^th^ residue. According to this finding, the drug binding may help to stabilize certain areas of the protein structure. The degree of protein compactness was looked at using the radius of gyration (Rg) study. Less folding was evident in the Rg (nm) values of the Levothyroxine-receptor complex compared to the apo receptor, suggesting that the compactness of the protein may have been impacted by the drug’s interaction. Lastly, Solvent Accessible Surface Area (SASA) analysis was used to forecast the stability of proteins’ hydrophobic cores. Initial SASA values for the apo receptor and drug-receptor complex were both 65 nm^2^, but after 90 ns, the values converged, showing that the drug’s interaction did not disrupt the protein’s hydrophobic core. Overall, the study’s findings point to the stability of the Levothyroxine-receptor complex and the possibility that the drug’s binding may have had an impact on the protein’s stability and structure, particularly in areas with high mobility. The development of more effective and targeted medications that target this receptor may be affected by these discoveries.

## Conclusion

In this study, 7 DEGs of AD were predicted and they could be targeted as potential biomarkers for the diagnosis of AD. These genes could be used to monitor disease progression and treatment response, leading to more personalized treatment options. One repurposable drug candidate, Levothyroxine was identified whose interaction with the target Transthyretin was confirmed through molecular docking and dynamics simulation analysis. Overall, the identification of potential biomarkers and therapeutics for AD could have significant implications for the diagnosis, treatment, and management of this incapacitating condition. However, *in vitro* and *in vivo* studies are necessary for further validation of our findings.

---

**Supplementary Table 1:** Significant GO and functional pathway enrichment analysis.

